# Structure-function relationship of *Gossypium hirsutum* NAC transcription factor, GhNAC4 with regard to ABA and abiotic stress responses

**DOI:** 10.1101/2020.05.28.121665

**Authors:** Vikas Shalibhadra Trishla, Pulugurtha Bharadwaja Kirti

## Abstract

Our previous study demonstrated that the expression of *GhNAC4*, a NAC transcription factor from cotton, was induced by abiotic stresses and abscisic acid (ABA) treatment. In the present study, we investigated the molecular mechanisms underlying ABA and stress response of GhNAC4. Over-expression of *GhNAC4* in transgenic tobacco conferred tolerance to salinity and drought treatments with associated enhanced expression of several stress-responsive marker genes. GhNAC4 is a nuclear protein that exhibits transcriptional activation property, and the C-terminal transcriptional regulatory (TR) domain is responsible for it. GhNAC4 also forms homo-dimers and the N-terminal NAC-domain is essential for this activity. The structure-function relationship of NAC transcription factors, particularly with respect to abiotic stress tolerance remains largely unclear. In this study, we investigated the domains essential for the biochemical functions of GhNAC4. We developed transgenic tobacco plants overexpressing the GhNAC4 NAC-domain and the TR-domain separately. NAC-domain transgenics showed hypersensitivity to exogenous ABA while TR-domain transgenics exhibited reduced sensitivity. Abiotic stress assays indicated that transgenic plants expressing both the domains separately were more tolerant than wild-type plants with the NAC-domain transgenics showing increased tolerance as compared to TR-domain transgenics. Expression analysis revealed that various stress-responsive genes were upregulated in both NAC-domain and TR-domain transgenics as compared to wild-type under salinity and drought treatments. These results suggest that the stress tolerance ability of GhNAC4 is associated with both the component domains while the ABA responsiveness is largely associated with N-terminal NAC-domain.

**Key Message:** NAC and transcriptional regulatory domains are responsible for the abiotic stress tolerance ability of the cotton NAC transcription factor GhNAC4 while the ABA-responsiveness is largely associated with the NAC-domain.

## Introduction

Transcription factors (TFs) are key modulators of gene expression patterns leading to the regulation of growth, development and environmental stress responses. A typical TF minimally consists of a DNA binding domain and a transcriptional regulatory domain. The earliest reports of the modular nature of TFs came from the ‘domain-swap’ experiment with the fusion of DNA binding domain of LexA (*Escherichia coli* repressor protein) and the activation domain of GAL4 TF (*Saccharomyces cerevisiae* transcriptional activator). This resulted in a chimeric transcriptional activator that identified LexA binding site (Brent and Ptashne 1985). This remarkable modular nature of TFs has been confirmed in many other systems (Graham et al. 1999; Hollenberg and Evans 1988; Porsch et al. 2005). The exact position of these domains/modules within the chimeric proteins is highly flexible suggesting that each domain represents an independent structural module with an independent function (Frankel and Kim 1991).

NAC (*NAM*, *ATAF1/2*, *CUC*) proteins constitute a prominent group of plant-specific TFs. Genomes of *Arabidopsis thaliana*, rice, tobacco and cotton contain more than 100 NAC TFs (Li et al. 2018; Nuruzzaman et al. 2010; Ooka et al. 2003; Sun et al. 2018). A typical NAC TF comprises a well-conserved N-terminal NAC-domain, which possesses DNA-binding and dimerization capability (Ernst et al. 2004) and a C-terminal transcription regulatory (TR) domain, which is highly variable for varied functions and is responsible for conferring transcriptional activation or repression function of NAC TFs (Olsen et al. 2005). NAC TFs play vital roles in diverse developmental and stress response processes in plants (Nakashima et al. 2007; Sablowski and Meyerowitz 1998; Souer et al. 1996; Sperotto et al. 2009).

Although several studies have investigated the function of NAC TFs, very few studies have aimed at dissecting the structure-function relationships of NAC TFs. Jensen et al. (2010) identified ANAC019 as a positive regulator of ABA responses and demonstrated that the ectopic expression of full length N-terminal NAC domain or C-terminal TR-domain of *ANAC019* alone conferred hypersensitivity to exogenous ABA. To understand the structure-function relationship of ANAC019 for ABA responsiveness, they developed chimeric proteins by swapping the NAC-domain and TR-domains with analogous regions from other NAC TFs and demonstrated that the biochemical and functional specificity of ANAC019 is associated with both the NAC and the TR-domains. In another study, Taoka et al. (2004) investigated the domain specificity of *Arabidopsis* CUC TFs, which play an essential role in shoot apical meristem formation. They demonstrated that TR-domains from three different NAC TFs could be fused to NAC-domain of CUC2 and could still retain the shoot induction functionality of CUC2. However, replacing the CUC2 NAC-domain with ATAF NAC-domain resulted in the loss of shoot induction functionality. Domain swapping of CUC2 TF revealed that the NAC-domain is essential for the adventitious shoot formation. However, the structure-function relationship of NAC TFs, particularly with respect to abiotic stress tolerance remains largely unclear.

We previously showed that the promoter region of *GhNAC4*, a NAC TF from cotton (*Gossypium hirsutum*) is induced by various phytohormones and environmental stress treatments. The *GhNAC4* promoter was also predicted to contain several phytohormone and stress related *cis*-acting elements that are involved in the regulation of gene expression. Studies of gene expression analysis of *GhNAC4* have also implicated its role in abiotic stress and ABA response (Trishla et al. 2020).

In the present study, we have observed that *GhNAC4* is a positive regulator of ABA and over-expression of *GhNAC4* shows increased abiotic stress tolerance. GhNAC4 is a nuclear protein, which possesses transcriptional activation property and is also able to form homo-dimers. To determine the domain essential for GhNAC4 function, we developed transgenic tobacco plants overexpressing the NAC-domain and the TR-domain of GhNAC4 independently and investigated their responses to ABA and stress treatments. The findings suggested that the ABA-responsiveness of GhNAC4 is largely associated with the NAC-domain while the abiotic stress tolerance functionality of GhNAC4 is associated with both NAC- and TR domains. These data provide new insights into the structure-function relationships of NAC TFs functionality with respect to abiotic stress tolerance.

## Material and Methods

### Plant material and growth conditions

Leaves from two weeks-old cotton plants (*Gossypium hirsutum* var. JK Durga) were used for isolation of full-length gene or the regions encoding NAC and TR-domains in *GhNAC4*. Leaves from *in vitro* grown two weeks-old tobacco (*Nicotiana tabacum* cultivar Samsun) were used for generation of transgenic plants expressing full length *GhNAC4* or the regions encoding individual domains. The plants were maintained in a growth chamber at 26 ± 2 °C with 16 /8 h of light/dark.

### Isolation and cloning of full-length CDS of *GhNAC4*

The full-length CDS of *GhNAC4* (80-1120, GenBank Accession no. EU706342.1), NAC-domain (80-496) and TRD (497-1120) were amplified using reverse transcribed cDNA as a template. The primers used are mentioned in Supplementary Table 1. The amplicons were cloned into pRT100 vector (GenBank no. A05521.1, Töpfer et al. 1987) for transcriptional fusion with the CaMV 35S promoter and polyadenylation signal. The expression cassettes were further cloned in to plant binary vector pCAMBIA 2300 (http://www.cambia.org) for constitutive expression of transgenes and *nptII* (kanamycin resistance) selection gene in transgenic tobacco.

### Tobacco transformation

Genetic Transformation of tobacco was performed using the leaf-disc method as described by Horsch et al. (1985) using *Agrobacterium tumefaciens* (strain EHA105). Transgenics plants were selected using kanamycin (150 mg/L) and homozygous T_2_ generation plants were used for various analyses. The primers used are mentioned in Supplementary Table 1. The constitutively expressing tobacco transgenics of full-length *GhNAC4*, NAC-domain and TR-domain were named as GhNAC4, GhNAC4-N, GhNAC4-C genotypes, respectively.

### Sub-cellular localization of GhNAC4

Full-length *GhNAC4* CDS was fused N-terminally to EGFP present in pEGAD vector (Cutler et al. 2000) at the *Sma*I and *Bam*HI sites. The pEGAD:*GhNAC4* fusion construct was transfected into onion (*Allium cepa*) epidermal peels by agroinfiltration as described by Xu et al. (2014). The peels were analyzed by laser scanning confocal microscopy (TCS SP2, Leica Microsystems, Germany). The experiment was performed in triplicates.

### Site-directed mutagenesis

Point mutations and deletion constructs of *GhNAC4* CDS were developed following the site-directed mutagenesis protocol of Edelheit et al. (2009). The primers for inducing various point mutations were designed with the help of PrimerX software (https://www.bioinformatics.org/primerx/).

### Transcription activation assay

Full-length *GhNAC4* CDS and a series of deletion derivatives were cloned separately into the GAL4 DNA-binding domain vector, pGBDU (James et al. 1996) using primers listed in Supplementary Table1. The constructs were transformed into the *Saccharomyces cerevisiae* strain pJ694a (James et al. 1996) using the polyethylene glycol-lithium acetate method (Gietz and Schiestl 2007) and plated on Synthetic Dropout (SD) medium lacking uracil (Ura^−^) for 3 d at 30 °C. For the qualitative growth assay, three colonies were picked and overnight grown cultures in SD Ura^−^ media were diluted to 1.0 OD units. They were then serially diluted 10-fold, and 5 μL was spotted on SD Ura^−^ media and SD Ura^−^ media either lacking histidine or adenine (Ura His^−^/ Ura Ade^−^). The transcriptional activity was evaluated based on the growth after 3 d at 30 °C. The empty vector pGBDU was used as a negative control. The experiment was repeated with three independent transformations. For the quantitative assay, β-galactosidase activity was measured by the O-nitrophenyl-β-galactopyranoside (ONPG) assay method.

### Yeast two-hybrid assay

Various deletion constructs of *GhNAC4* CDS were cloned into the GAL4 DNA-binding domain (pGBDU) and the activation domain (pGAD) vectors (James et al. 1996) using primers listed in Supplementary Table1. The constructs were transformed into the yeast strain, pJ694a and plated on SD medium lacking uracil and leucine (Ura Leu^−^) for 5 d at 30 °C. For the qualitative growth assay, three colonies were picked and overnight grown cultures in SD Ura Leu^−^ media. They were then serially diluted 10-fold, and 5 μL was spotted on SD Ura Leu^−^ media and SD Ura Leu^−^ media either lacking histidine or adenine (Ura Leu His^−^/ Ura Leu Ade^−^). The protein-protein interaction was evaluated based on the growth after 5 d at 30 °C. The empty vectors pGBDU and pGAD were used as negative controls. The experiment was repeated with three independent transformations. For the quantitative assay, β-galactosidase activity was measured by the ONPG assay method.

### β-galactosidase assay

β-galactosidase activity was essentially assayed following Miller (1972) with the following modifications. Single yeast colony was grown in selective media (Ura^−^ or Ura Leu^−^) overnight, and secondary cultures were grown to an absorbance (OD_600_) of 0.8-1.0. A 5 mL sample of the secondary culture was pelleted by centrifugation at 2,000 × g and resuspended in 5 mL of Z-buffer and OD_600_ was again recorded. The composition of the Z-buffer is 60 mM Na2HPO4, 40 mM NaH2PO4, 10 mM KCl, 1 mM MgSO4 and 50 mM β-mercaptoethanol pH 7.0. To an 800 μL aliquot of the culture, 50 μl each of 0.1% sodium dodecyl sulphate and chloroform was added and the mixture was vortexed briefly for permeabilization. After 5 min incubation, 160 μL of ONPG solution (4 mg/mL in Z-buffer, HiMedia) was added and vortexed briefly before incubating at 30 °C for 2 min to 4 h. The reaction was terminated by the addition of 400 μL of 1 M Na_2_CO_3_. The sample was centrifuged for 2 min at 16000 × g, and the absorbance of the supernatant read at 420 nm and 550 nm. The enzyme activity was expressed in Miller units. Mean values and standard error were calculated from four measurements obtained from three independent yeast transformants.

### Seed germination assay

For the seed germination assay, seeds of WT, GhNAC4, GhNAC4-N, GhNAC4-C genotypes were harvested at the same time and stored under the same conditions. They were surface-sterilized in 70% (v/v) ethanol for 2 min, followed by 15 min in 4% (w/v) in aqueous sodium hypochlorite and rinsed five times with sterile distilled water. Seeds were sown on half-strength salts of Murashige-Skoog (MS) media (Murashige and Skoog 1962) containing 0.6% Gelzan (Sigma-Aldrich) pH 5.7, supplemented with or without 150 mM NaCl and 2 μM ABA (mixed isomers, Sigma-Aldrich). The −0.5 MPa PEG 8000 (HiMedia) infused plates were prepared according to van der Weele et al (2000). Germination rates were scored on the 7 d for radicle emergence or presence of green cotyledons. The experiment was conducted in triplicate with 80-100 seeds for each genotype and experiment.

### Root elongation assay

For the post-seed germination growth assay, 7 days old seedlings germinated on half-strength MS media were transferred to media containing 200 mM NaCl and 10 μM ABA (mixed isomers, Sigma-Aldrich). The −0.7 MPa PEG 8000 (HiMedia) infused plates were prepared according to van der Weele et al (2000). All the plates were placed upright in the growth chamber. The primary root lengths were measured on 15 d post-transfer. The experiment was conducted in triplicate with at least 18-21 seedlings for each genotype and experiment.

### Stomatal bioassay

Epidermal strip assay was performed as described by McAinsh et al (1996) with slight modifications. Epidermal strips from the abaxial side were peeled and floated in opening buffer (10 mM MES-KOH and 50 mM KCl, pH 6.1) under a photon flux density of 200 μmol m^2^ s^−1^ for 3 h. Subsequently, 10 μM ABA (dissolved in 95% ethanol) and an equal volume of ethanol (used as control) were added to the buffer. After 150 min of incubation, the peels were imaged using the light microscope (CX21, Olympus). The width of the stomatal aperture was measured using the pre-calibrated ocular micrometer of ProgRes CapturePro image analyzer (Jenoptik, Germany). Approximately, 60 stomatal apertures were measured for each genotype and the experiment was repeated three times.

### Water loss assay

Leaves from well-watered one-month-old WT, GhNAC4, GhNAC4-N and GhNAC4-C plants were excised and weighed immediately. The excised leaves were left at room temperature and weighed every 1 h (Li et al. 2019). Water loss was calculated as a percentage of loss in fresh weight over a period of time. Six leaves from separate plants for each genotype were used and the experiment was repeated three times.

### Leaf disc assay

The third leaf from the top of one-month-old plants of WT, GhNAC4, GhNAC4-N and GhNAC4-C genotypes were used for the leaf disc assay. Using a cork borer, leaf discs of 1 cm diameter were punched and floated on 200 mM NaCl and 15 % PEG 8000 for 4 d at 24 ± 2 °C. The experiment was repeated three times.

### Total chlorophyll content

The 100 mg samples of treated leaf discs were used for extracting total chlorophyll using 80 % acetone according to Arnon (1949). Total chlorophyll content was expressed as mg/g FW. This experiment was repeated three times.

### The extent of lipid peroxidation

The treated leaf discs (100 mg) were homogenized in 1 ml of 0.1 % trichloroacetic acid (TCA) and centrifuged at 10,000 × g for 10 min. To the supernatant, 5 ml of thiobarbituric acid (0.5 % in 20 % TCA solution) was added and the mixture was boiled for 25 min at 100 °C. The reaction was terminated by quick cooling on ice, followed by centrifugation at 10,000 × g for 5 min. The supernatant was used to measure malondialdehyde (MDA) content according to Heath and Packer (1968). The MDA content was expressed as nmoles/g FW. The experiment was repeated three times.

### Proline content

The treated leaf discs (100 mg) were homogenized in 0.5 ml of 3 % sulfosalicylic acid and centrifuged at 10,000 × g for 10 min. To 100 μl of supernatant, a mixture of 100 μl of 3 % sulfosalicylic acid, 200 μl of glacial acetic acid and 200 μl of acid ninhydrin were added. The mixture was boiled for 60 min at 100 °C and the reaction was terminated by quick cooling on ice. The samples were extracted with toluene and the upper organic phase was used for measuring the proline content according to Bates (1973). The proline content was expressed as μmoles/g FW. The experiment was repeated three times.

### Expression analysis of stress marker genes in tobacco

To assess the transcript levels of ABA and stress-responsive genes in tobacco seedlings, quantitative real-time PCR (qPCR) was employed. For this, two weeks old seedlings of wild-type, GhNAC4, GhNAC4-N, GhNAC4-C genotypes were treated with 200 mM NaCl and 15% PEG 8000 for 24 h. Total RNA was extracted using Trizol reagent (Invitrogen, USA) and DNA contamination was removed by treating it with RNase-free DNase I (Takara Bio, China). Approximately 1 μg of total RNA was used for the first-strand cDNA synthesis using RevertAid cDNA synthesis kit (Thermo Fischer Scientific, USA) according to the manufacturer’s instructions. For the qPCR, cDNA was diluted to 100 ng/μl and was mixed with Green Premix Ex Taq II (Takara Bio, China) and amplifications were carried out following the manufacturer’s protocol. The constitutively expressing Ubiquitin gene (*NtUBI1*, GenBank Accession no. U66264.1) from tobacco was used as an internal reference gene. The stress-responsive genes used in the study are *NtAPX* (U15933.1), *NtCAT1* (U93244.1), *NtERD10C* (AB049337.1), *NtERF5* (AY655738.1), *NtDREB3* (EU727157.1), *NtMnSOD* (AB093097. 1), *NtNCED3* (JX101472.1), *NtSOS1* (XM_009789739.1), and *NtSUSY* (AB055497.1). The primers of stress-associated genes are mentioned in Supplementary Table 2. The fold change was determined using the ΔΔC_T_ method (Livak and Schmittgen 2001). The experiments were performed in triplicates, and two independent biological replicates were used for the analyses.

### Statistical analysis

Data were analyzed by one-way analysis of variance (ANOVA) using SigmaPlot 11.0 software. The data were expressed as the mean ± SE. \*\*\**P* < 0.001, \*\**P* < 0.01 and \**P* < 0.05 represent significant differences at 0.1, 1 and 5% level respectively.

## Results

In our previous study, we identified that *GhNAC4* is highly induced in PEG, NaCl and ABA treatments in cotton (Figs. 1 and 2, Trishla et al. 2020). This has prompted us to further examine the role of GhNAC4 protein in abiotic stress responses.

**Fig. 1.**
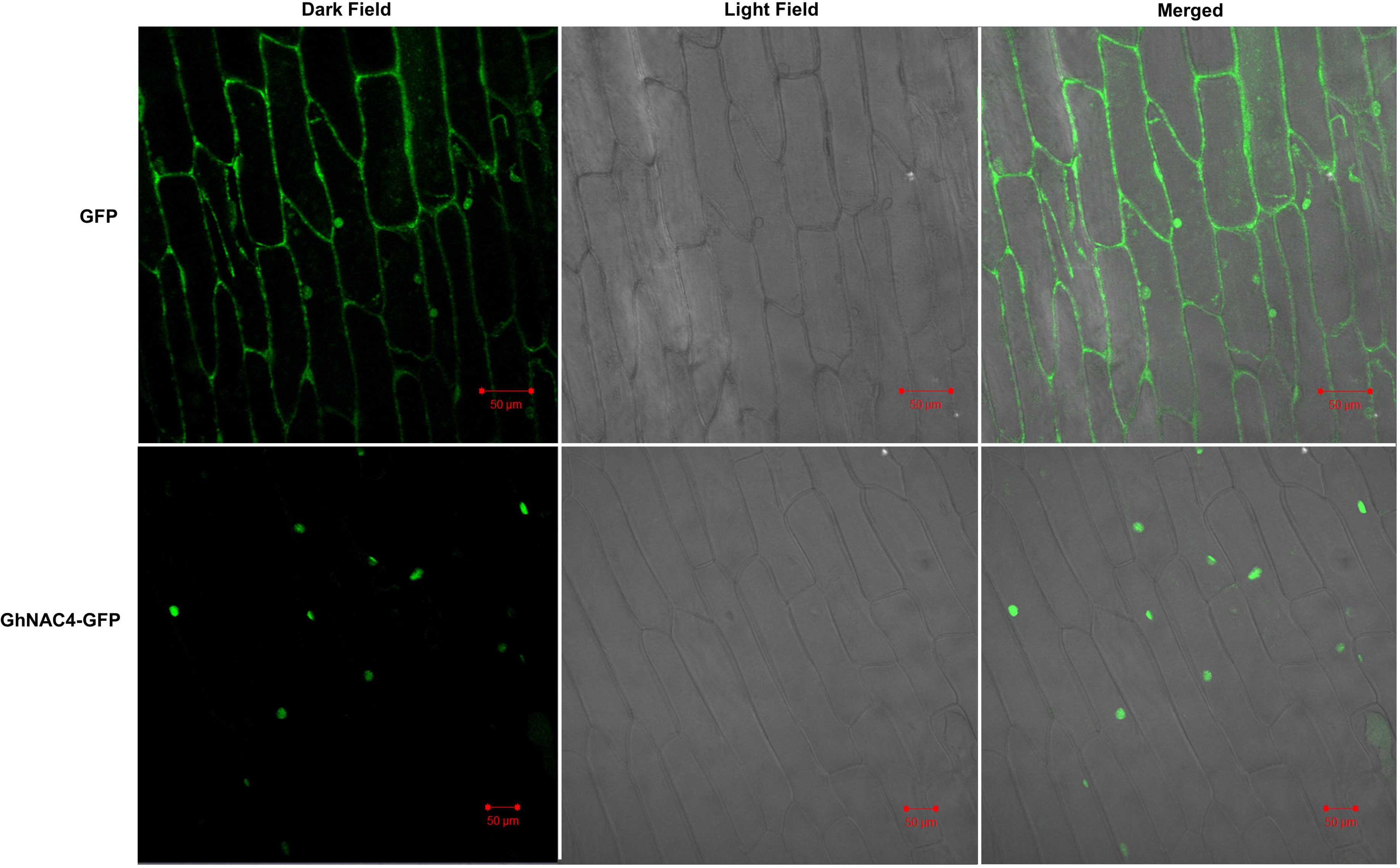
Sub-cellular localization of GhNAC4. Empty pEGAD vector and pEGAD:*GhNAC4* recombinant vectors were agroinfiltrated in onion epidermal peels for transient expression. The fluorescence was analyzed by laser scanning confocal microscopy. The experiment was repeated three times and representative images have been presented here. Scale bar = 50 μM

**Fig. 2.**
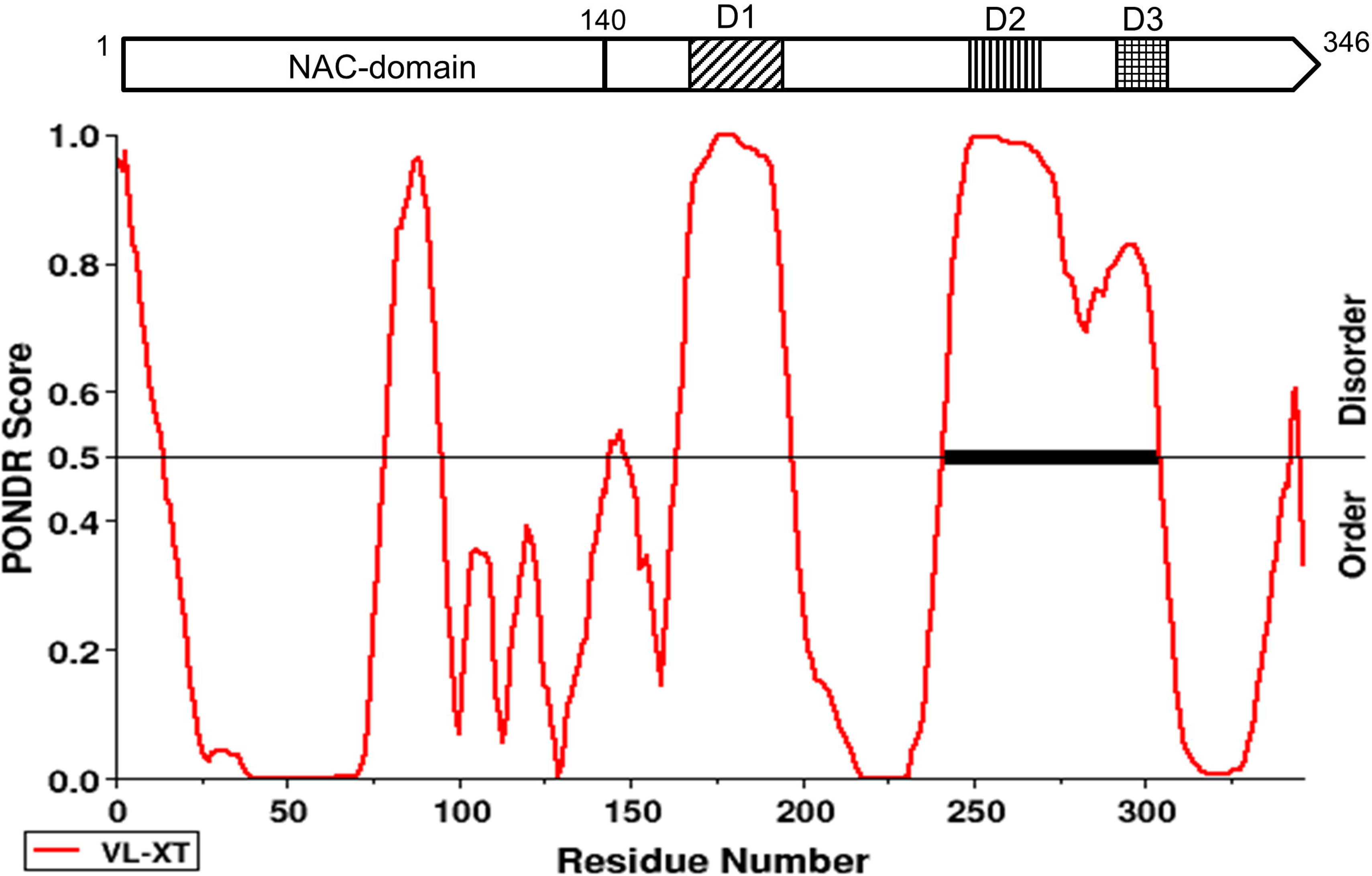
Structure of GhNAC4. Top, Pictorial representation of the predicted structure of GhNAC4 drawn to scale with N-terminal NAC domain and the C-terminal showing three intrinsically disordered regions, D1, D2 and D3. Bottom, prediction of intrinsically disordered regions of GhNAC4 using PONDR (http://www.pondr.com/). The threshold for disordered regions is ≥ 0.5

### GhNAC4 is a nuclear protein

The recombinant construct carrying the pEGAD:*GhNAC4* translational fusion was agroinfiltrated in onion epidermal cells and visualized through laser scanning confocal microscopy. The GhNAC4:EGFP fusion protein has been identified to localize in the nucleus (Fig. 1). The EGFP of the empty pEGAD vector, which was used as a control was observed throughout the whole cell. This result suggests that GhNAC4 is a nuclear-localized protein.

### GhNAC4 possesses transcription activation property

GhNAC4 has the structure of a typical NAC TF and is predicted to possess an N-terminal NAC-domain and a C-terminal intrinsically disordered transcription regulatory domain (Fig. 2). To verify whether GhNAC4 possessed transcription activation property, it was fused to the GAL4 DNA binding domain, as shown in Fig. 3. The transcription activation was examined by growing the transformants on Ura^−^, Ura His^−^ and Ura Ade^−^ media and quantifying the β-galactosidase activity. The full-length GhNAC4 (1-346 aa) and the C-terminal truncated variant, GhNAC4Δ140-346-C showed growth on all media whereas the N-terminal truncated variant, GhNAC4Δ1-139-N failed to grow. It suggests that GhNAC4 is a transcription activator, and the C-terminal region of the protein is responsible for this property. Interestingly, the β-gal assay indicates that the C-terminal variant possesses twice the activity as compared to the full-length protein.

**Fig. 3.**
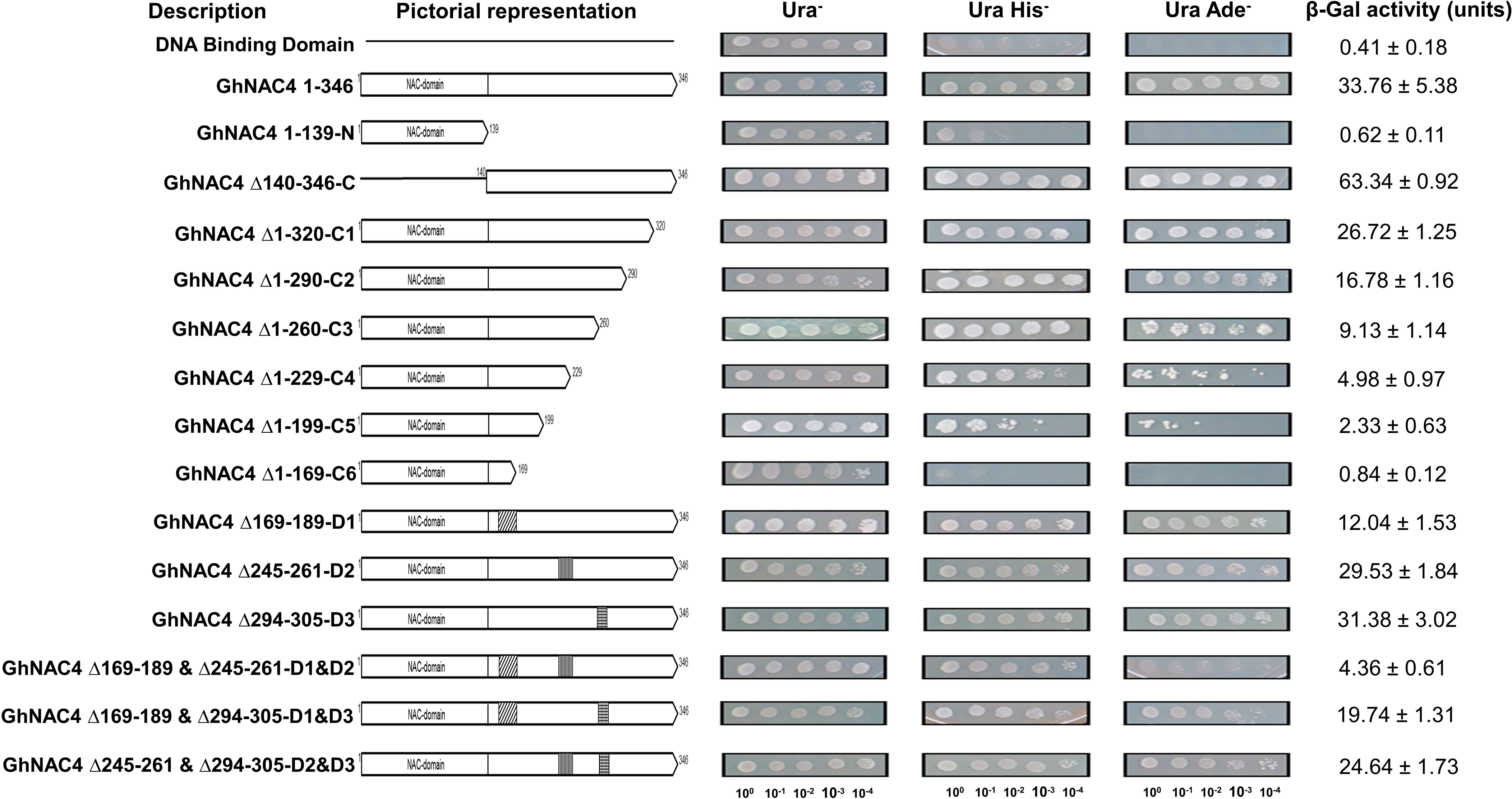
Transcription activation of GhNAC4 in yeast and mapping of the activation domain. Various deletion constructs of *GhNAC4* (left) were fused to the GAL4 DNA-binding domain of pGBDU vector and transformed in yeast strain, pJ694a and screened for transcription activation assay. Transformed yeast cells were serially diluted 10-fold and spotted (right) onto plates lacking uracil (Ura^−^) to check for transformation and on plates lacking uracil and histidine (Ura His^−^) and uracil and adenine (Ura Ade^−^) to allow the screening of weak and strong transcription activation activity respectively. Empty pGBDU vector was used as a negative control. β-galactosidase activity (units), represented as mean ± SE from three independent transformations, is indicative of transcription activation (extreme right)

To determine the minimum region essential for the transcription activation property, a progressive series of deletion variants of GhNAC4 were analyzed in yeast. The minimal truncated region that showed transcription activation was from amino acid 140-199 (GhNAC4Δ1-199-C5).

Further deletion completely abolished the transcriptional activation (GhNAC4Δ1-169-C6) property. The β-gal assay demonstrated that progressive deletion of 30 amino acids reduced the activity by half, as indicated in Fig. 3.

GhNAC4 C-terminal region is predicted to possess three intrinsically disordered regions (Fig. 2). To determine the region of the C-terminal domain necessary for the transcription activation property, we made stepwise and combination deletions of these low-complexity regions (Fig. 2). Removal of the regions GhNAC4Δ169-189-D1 and GhNAC4Δ245-261-D2 in combination almost abolished the transcription activation. This indicates the importance and dependency of the transcription activation property on the intrinsically disordered regions.

### GhNAC4 exists as a dimer

Ernst et al. (2004) reported that ANAC019 exists as a dimer. The formation of two salt bridges by the conserved Arginine 19 and Glutamate 26 are among the prominent interactions between the two dimers. Strong transcription activation capability of the full-length GhNAC4 and the C-terminal domain (GhNAC4Δ140-346-C) prevents protein-protein interaction studies. To assess the homo-dimerization property of GhNAC4, a yeast two-hybrid assay was carried out using the N-terminal NAC-domain (GhNAC4Δ1-139-N), as shown in Fig. 4. The protein-protein interaction was examined by growing the transformants on Ura Leu^−^, Ura Leu His^−^ and Ura Leu Ade^−^ media and quantifying the β-galactosidase activity. The N-terminal domain was able to homo-dimerize although weakly. Any point mutation to disrupt either one or both the salt bridges resulted in the complete abolition of dimerization as shown in Fig. 4. This indicates the importance and dependency of the salt bridges for the homo-dimerization property of GhNAC4 protein.

**Fig. 4.**
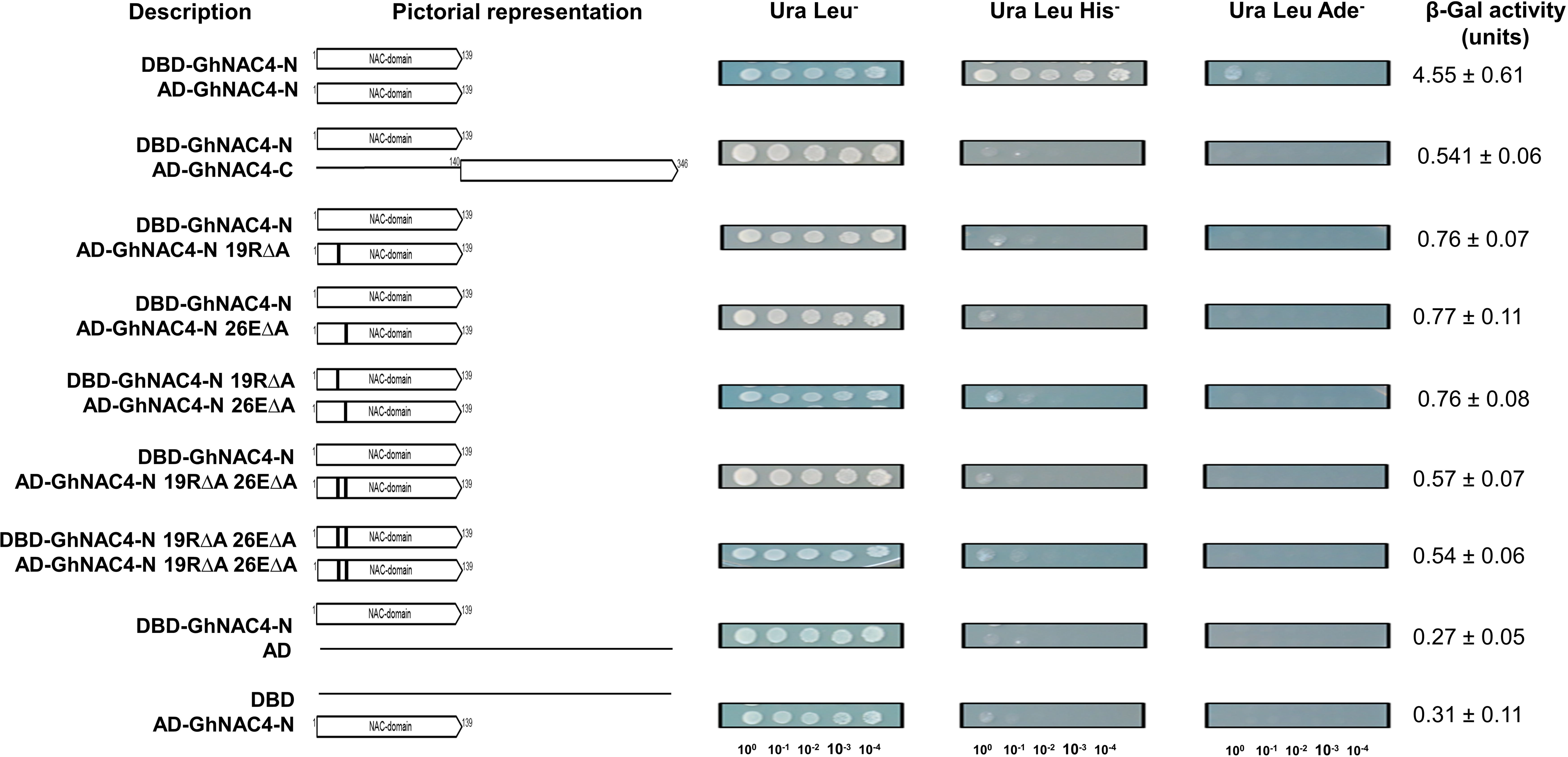
Homo-dimerization of GhNAC4. Various deletion constructs of *GhNAC4* (left) were fused to the GAL4 DNA-binding domain (DBD) of pGBDU vector and GAL4 activation domain (AD) of pGAD vector and transformed in yeast strain, pJ694a to screen for homo-dimerization assay. Transformed yeast cells were serially diluted 10-fold and spotted (right) onto plates lacking uracil and leucine (Ura Leu^−^) to check for transformation and on plates lacking uracil, leucine and histidine (Ura Leu His^−^) and uracil, leucine and adenine (Ura Leu Ade^−^) to allow the screening of weak and strong protein-protein interaction activity, respectively. Empty pGBDU and pGAD vectors were used as negative controls. β-galactosidase activity (units), represented as mean ± SE from three independent transformations, is indicative of protein-protein interaction (extreme right)

### The NAC-domain is sensitive to ABA-mediated seed germination and post-germinative growth and both domains are required for stress tolerance

The increased expression of *GhNAC4* in response to exogenous ABA, NaCl and PEG (Figs. 1 and 2, Trishla et al. 2020) suggested that GhNAC4 might play a role in ABA signaling and abiotic stress responses. ABA plays a crucial role in abiotic stress response as well as in seed germination and seedling growth (Nakashima and Yamaguchi-Shinozaki 2013). Thus, to investigate the role of the full-length, NAC- and the TR-domains of GhNAC4 protein in ABA and abiotic stress responses, we generated transgenic tobacco plants over-expressing GhNAC4 and its domains independently (Supplementary Figure 1). We examined the germination rates of wild type, GhNAC4, GhNAC4-N, GhNAC4-C T_2_ generation seeds under various conditions as shown in Fig. 5. Seeds of these genotypes, germinated on ½ MS media showed no significant differences in germination rate under normal growth conditions. However, the germination of all genotypes under salinity stress (150 mM NaCl) was severely affected (Fig. 5). Wild-type showed less than 15% germination frequency while GhNAC4-C genotype showed around 28%. Interestingly, the germination rate of GhNAC4 and GhNAC4-N genotypes reached around 45% and 38% respectively, seven days after sowing. Drought stress-induced by PEG (−0.5 MPa) resulted in decreased germination frequency of wild-type and GhNAC4-C genotypes by 20% and 40% respectively. However, the germination frequency still reached above 70% for both the GhNAC4 and GhNAC4-N genotypes, seven days after sowing (Fig. 5). This indicates that the stress response of GhNAC4 is associated with both NAC-domain and TR-domain.

**Fig. 5.**
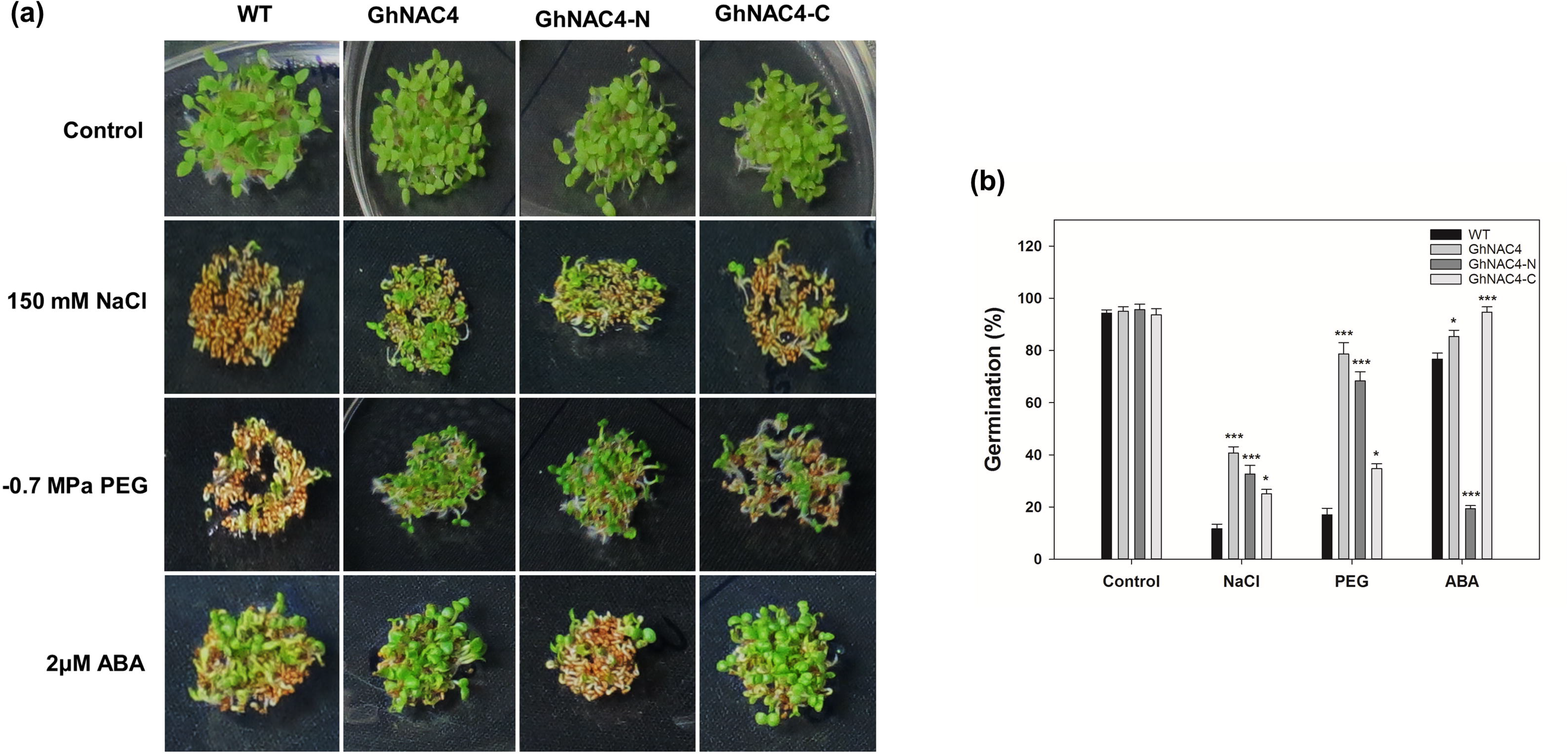
Responses of WT, GhNAC4, GhNAC4-N and GhNAC4-C expressing tobacco transgenic plants to NaCl and PEG induced stress at seed germination stage. **a** Representative photograph of germination of seeds on the 7 d, sown on control medium (half-strength MS) or media containing 150 mM NaCl, −0.5MPa PEG 8000 or 2 μM ABA. **b** Quantification of the percentage of germinated seeds of each genotype on the 7 day on the indicated treatments as described in **a**. For each experiment, approximately 80-100 seeds were scored per genotype. Data are means ± SE (n=3). A statistical analysis with one-way ANOVA indicates significant differences (*** *P* < 0.001 and * *P* < 0.05)

We next determined the ABA sensitivity of the wild type, GhNAC4, GhNAC4-N, GhNAC4-C genotype during germination. Treatment with 2 μM ABA severely arrested the growth of GhNAC4-N genotype seeds after the emergence of the radicle suggesting hypersensitivity. Interestingly, GhNAC4-C showed very little inhibition with germination frequency reaching over 95% suggesting near insensitivity. The germination frequencies of wild-type and GhNAC4 genotypes were also inhibited to a lesser extent (Fig. 5). This indicates that the NAC-domain is mainly responsible for the ABA receptivity of GhNAC4.

Subsequently, wild-type, GhNAC4, GhNAC4-N, GhNAC4-C genotype seedlings were assessed for their responses to abiotic stresses and ABA during the post-germinative growth stage as shown in Fig 6. Seedlings of these genotypes showed no significant differences in root length under normal growth conditions. However, under salinity stress (200 mM NaCl), the growth of all genotypes was severely affected (Fig. 6). Root length in the wild-type reduced to approximately 18%, while GhNAC4-C genotype around 28%. The primary root length of GhNAC4 and GhNAC4-N genotypes reached around 77% and 60% respectively. In addition, drought-induced by PEG (−0.7 MPa) also caused a reduction in root length of GhANC4, GhNAC4-N and GhNAC4-C genotypes to approximately 90%, 67% and 40% respectively whereas the wild-type seedlings root length reduced to 19% (Fig. 6). This result clearly indicated that GhNAC4 overexpressing plants are tolerant to salinity and drought stress and this property is conferred by both NAC- and TR-domains.

**Fig. 6.**
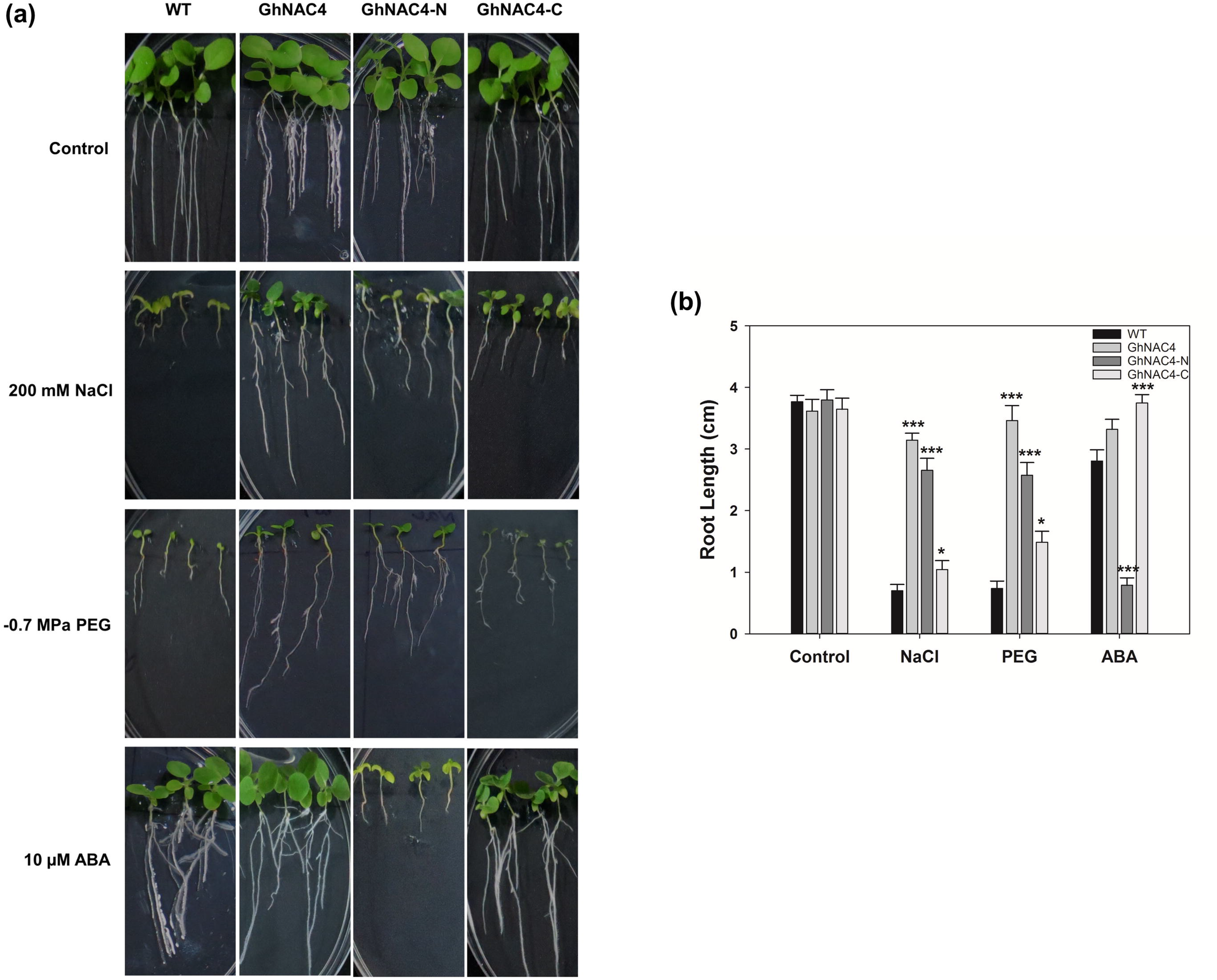
Responses of WT, GhNAC4, GhNAC4-N and GhNAC4-C expressing tobacco transgenic plants to NaCl and PEG induced stress at post-germination stage. **a** Representative photograph of seedlings at 15 d after transfer to control medium (half-strength MS) or media containing 200 mM NaCl, −0.7MPa PEG 8000 or 10 μM ABA. Seeds were germinated on half-strength MS for 7 d before transfer. **b** Quantification of root lengths for seedlings treated as described in **a**. For each experiment, approximately 18-21 seedlings were scored per genotype. Data are means ± SE (n=3). A statistical analysis with one-way ANOVA indicates significant differences (*** *P* < 0.001 and * *P* < 0.05)

Next, we considered the possible effect of ABA on the root growth. As in the case of seed germination, the GhNAC4-N seedlings showed hypersensitivity to ABA (10 μM) with significantly lower primary root growth (approximately 20%) and whereas the GhNAC4-C seedlings were insensitive (Fig. 6). This further confirms the ABA receptivity role of NAC-domain in GhNAC4.

### NAC-domain transgenics show enhanced stomatal closure under ABA treatment

To understand the physiological roles of the NAC and TR-domains, we investigated the behaviour of guard cells under ABA treatment. As seen in Fig. 7 stomatal aperture of GhNAC4-N and GhNAC4-C genotypes, respectively were ~ 20 and 36% smaller that than that of wild-type plants, under control conditions while there was no significant difference in the stomatal aperture of GhNAC4 plants. ABA is known to induce stomatal closure (Bright et al. 2006) and treatment with 10 μM ABA promoted stomatal closure in all genotypes (Fig. 7). However, the leaves of GhNAC4-N genotype showed a greater percentage of stomatal closure (~ 60%). In GhNAC4-C plants, the same treatment reduced the stomatal aperture by ~ 17%. Wild type and GhNAC4 plants showed a reduction by ~ 40-46% (Fig. 7). This corroborated the observation that GhNAC4-N genotype shows hypersensitivity to ABA while GhNAC4-C genotype is nearly insensitive and GhNAC4 is a positive regulator of ABA-mediated stomatal closure.

**Fig. 7.**
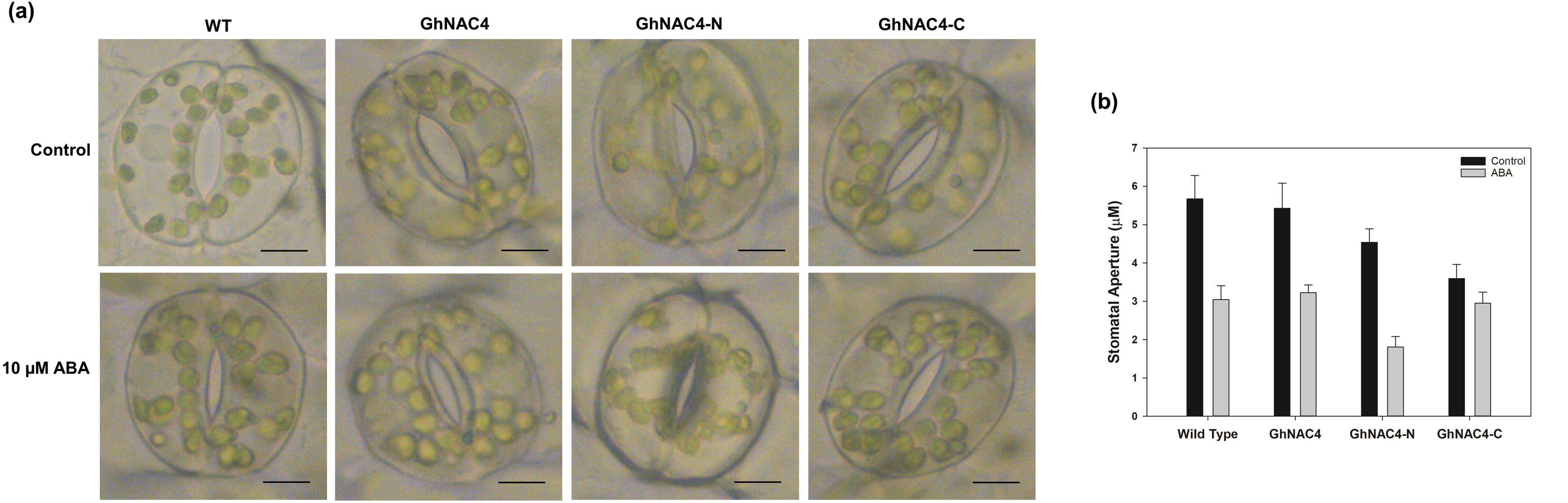
Responses of WT, GhNAC4, GhNAC4-N and GhNAC4-C expressing tobacco transgenic plants to ABA-induced stomatal closure. Epidermal peels prepared from abaxial sides of the leaves were floated in MES-KCl buffer for 3 h under high-light conditions to induce full opening of stomata. Subsequently, they were treated with 10μM ABA for 150 min. **a** A representative image from each genotype is presented. Scale bar = 5μm. **b** Quantification of stomatal aperture treated as described in **a**. Following treatment, the width of the stomatal aperture was measured. The experiment was performed in triplicate with 60 pairs of guard cells per genotype per experiment. Values are means ± SE

The modulation of the NAC and TR-domains in ABA-mediated stomatal closure in the transgenic plants has again prompted us to examine whether they show differences in the regulation of dehydration responses. To investigate this, we excised leaves of wild-type, GhNAC4, GhNAC4-N and GhNAC4-C genotypes and water loss was examined over a period of time (Fig 8). After dehydration for 9 h, the water loss from excised leaves was higher for wild-type plants (~ 75%) as compared to the GhNAC4 plants (~ 25%). The excised leaves of GhNAC4-N genotype lost water more slowly (~ 35%) as compared to GhNAC4-C genotype (~ 55%) (Fig. 8). An increase in water loss of GhNAC4-C plants could be partially due to the insensitivity to ABA-mediated stomatal closure as compared to GhNAC4-N plants.

**Fig. 8.**
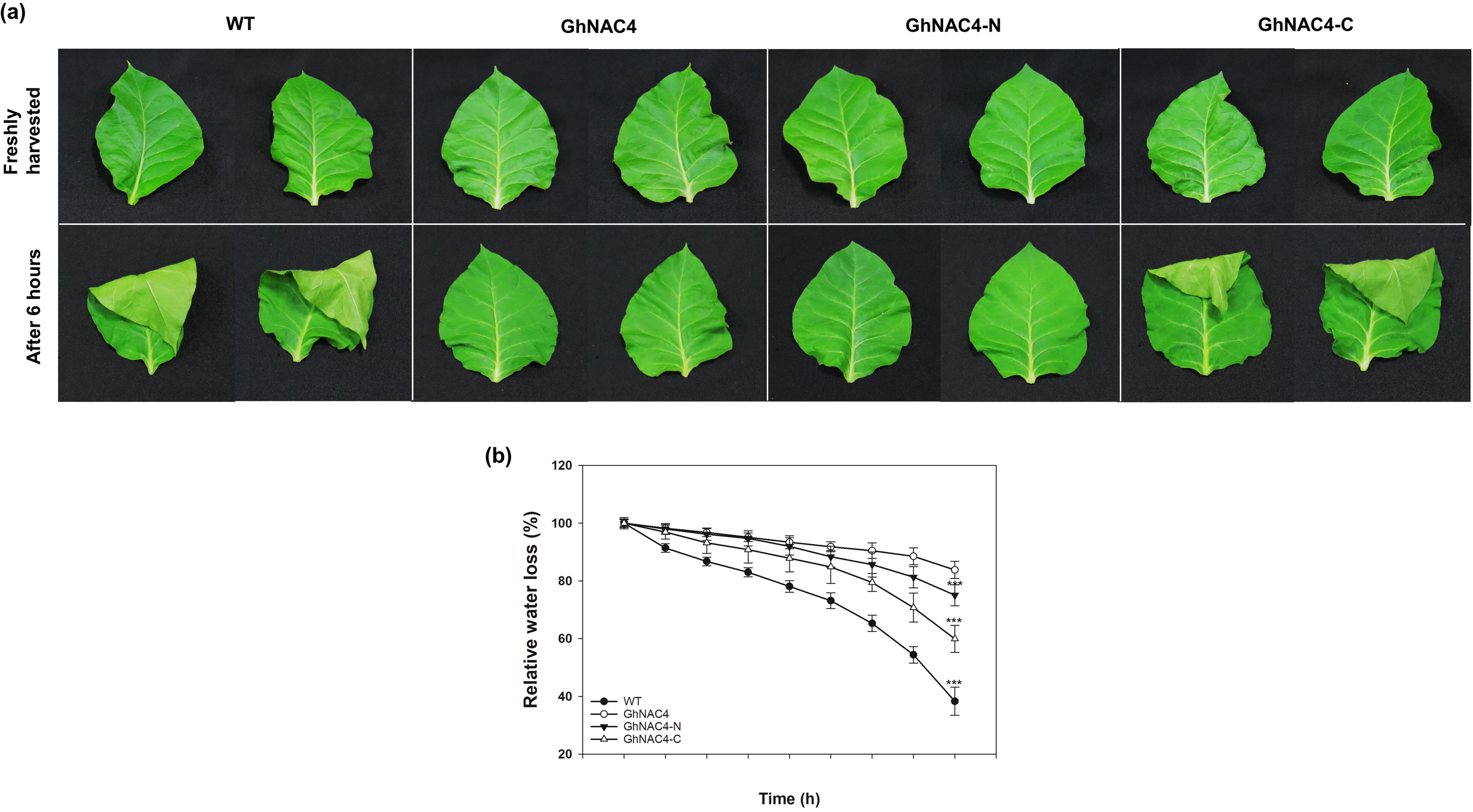
Water loss from WT, GhNAC4, GhNAC4-N and GhNAC4-C expressing tobacco transgenic plants. Leaves from one-month old plants were excised, immediately weighed and kept at room temperature. To measure water loss, fresh weight was weighed at regular intervals. **a** Photograph was taken at 6 h. A representative image from each genotype is presented. **b** Six leaves from separate plants for each genotype were used and the experiment was performed in triplicate. Values are means ± SE. A statistical analysis with one-way ANOVA indicates significant differences (*** *P* < 0.001)

### Both NAC and TR-domains are required for stress tolerance during vegetative growth

To assess the stress tolerance capabilities of wild-type, GhNAC4, GhNAC4-N and GhNAC4-C genotypes, leaf discs from fully expanded leaves were floated on 200 mM NaCl and 15 % PEG solutions (Fig. 9a). After 4 d of incubation, we observed a significant amount of bleaching in the leaf discs of wild-type and GhNAC4-C genotypes as compared to GhNAC4 and GhNAC4-N in both treatments. However, there was no significant difference in the chlorophyll content of all the corresponding leaf discs under control conditions. Under NaCl treatment, leaf discs of wild-type and GhNAC4-C demonstrated ~ 48-53 % bleaching. The total amount of chlorophyll lost from the leaf discs of GhNAC4 and GhNAC4-N was ~ 24-28 % under similar conditions. Dehydration stress caused by PEG treatment resulted in ~ 24-26 % bleaching in the leaf discs of wild-type and GhNAC4-C as compared to GhNAC4 and GhNAC4-N genotypes (~ 6-14 %) as seen in Fig. 9b.

**Fig. 9.**
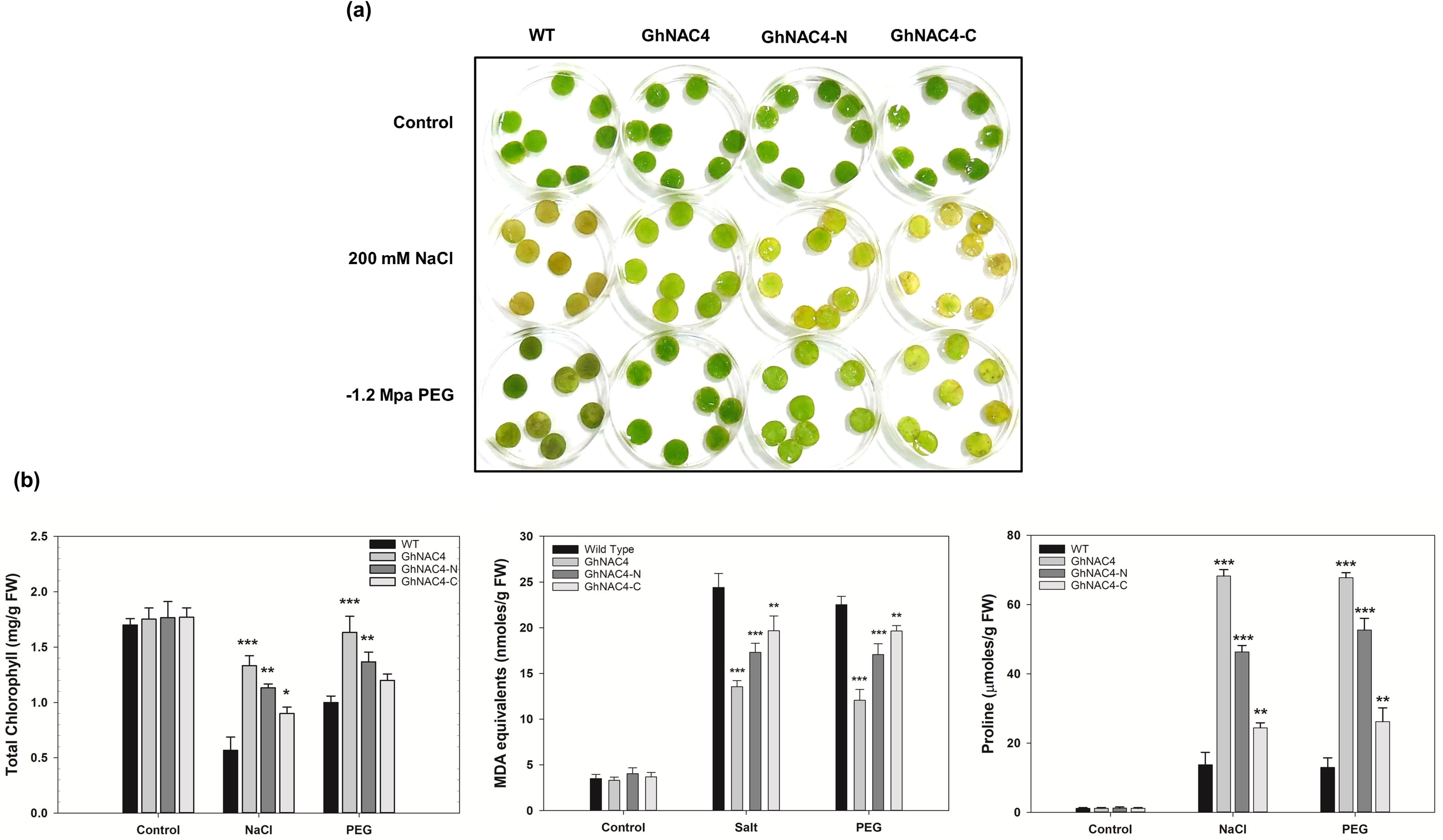
Response of WT, GhNAC4, GhNAC4-N and GhNAC4-C expressing tobacco transgenic plants to NaCl and PEG induced stress during vegetative growth. **a** Phenotype of leaf discs of wild type, GhNAC4, GhNAC4-N and GhNAC4-Cplants floated on 200 mM NaCl and 15 % PEG 8000 for 4 d. The experiment was repeated three times and representative photograph has been presented here. **b** The leaf discs were analyzed for the effect of NaCl and PEG induced stress on total chlorophyll content, extent of lipid peroxidation and proline accumulation. Values are mean ± SE (n=3). A statistical analysis with one-way ANOVA indicates significant differences (*** *P* < 0.001, ** *P* < 0.01 and * *P* < 0.05)

Salinity and dehydration treatments induce lipid peroxidation as a consequence of increased osmotic stress (Golldack et al. 2014). This is measured by malondialdehyde (MDA) content using the TBARS assay. There was an increase in the MDA content in all the genotypes under both the treatments. However, GhNAC4 and GhNAC4-N recorded a significantly lower content of MDA as compared to wild-type and GhNAC4-C (Fig. 9b).

Under abiotic stress conditions, plants produce osmolytes such as proline to maintain the redox balance (Szabados and Savouré 2010). Hence, proline content was measured in leaf discs of wild-type, GhNAC4, GhNAC4-N and GhNAC4-C genotypes floated on salt and PEG solutions. As compared to wild-type, GhNAC4 had a significantly higher amount of proline. Under similar conditions, GhNAC4-N recorded a significantly higher amount of proline as compared to GhNAC4-C (Fig. 9b). The above data allowed us to conclude that GhNAC4 is a positive regulator of salinity and drought stress tolerance. Further, both the NAC and TR-domains are likely to be involved in the adaptive processes of GhNAC4. However, the NAC-domain plays a more significant role in this regard.

### Both NAC and TR-domains regulate the expression of stress-responsive genes

Given the differential role NAC and TR-domains in ABA and abiotic stress responsiveness, it is pertinent to question whether they alter the expression of abiotic stress and ABA-responsive genes. We analyzed the expression of nine stress-responsive genes – *NtAPX* (ascorbate peroxidase), *NtCAT1* (catalase 1), *NtERD10C* (early responsive to dehydration 10C), *NtERF5* (ethylene response factor 5), *NtDREB3* (dehydration-responsive element-binding 3), *NtMnSOD* (superoxide desmutase), *NtNCED3* (9-*cis-*epoxycarotenoid dioxygenase), *NtSOS1* (salt overly sensitive 1), and *NtSUSY* (sucrose synthase) under NaCl and PEG treatments in all the genotypes. Under control conditions, the expression levels of all the 9 genes were very low and no significant differences were observed between wild-type, GhNAC4, GhNAC4-N, GhNAC4-C genotypes. However, under NaCl and PEG treatment, the transcript levels of all the nine genes were up-regulated (Fig. 10).

**Fig. 10.**
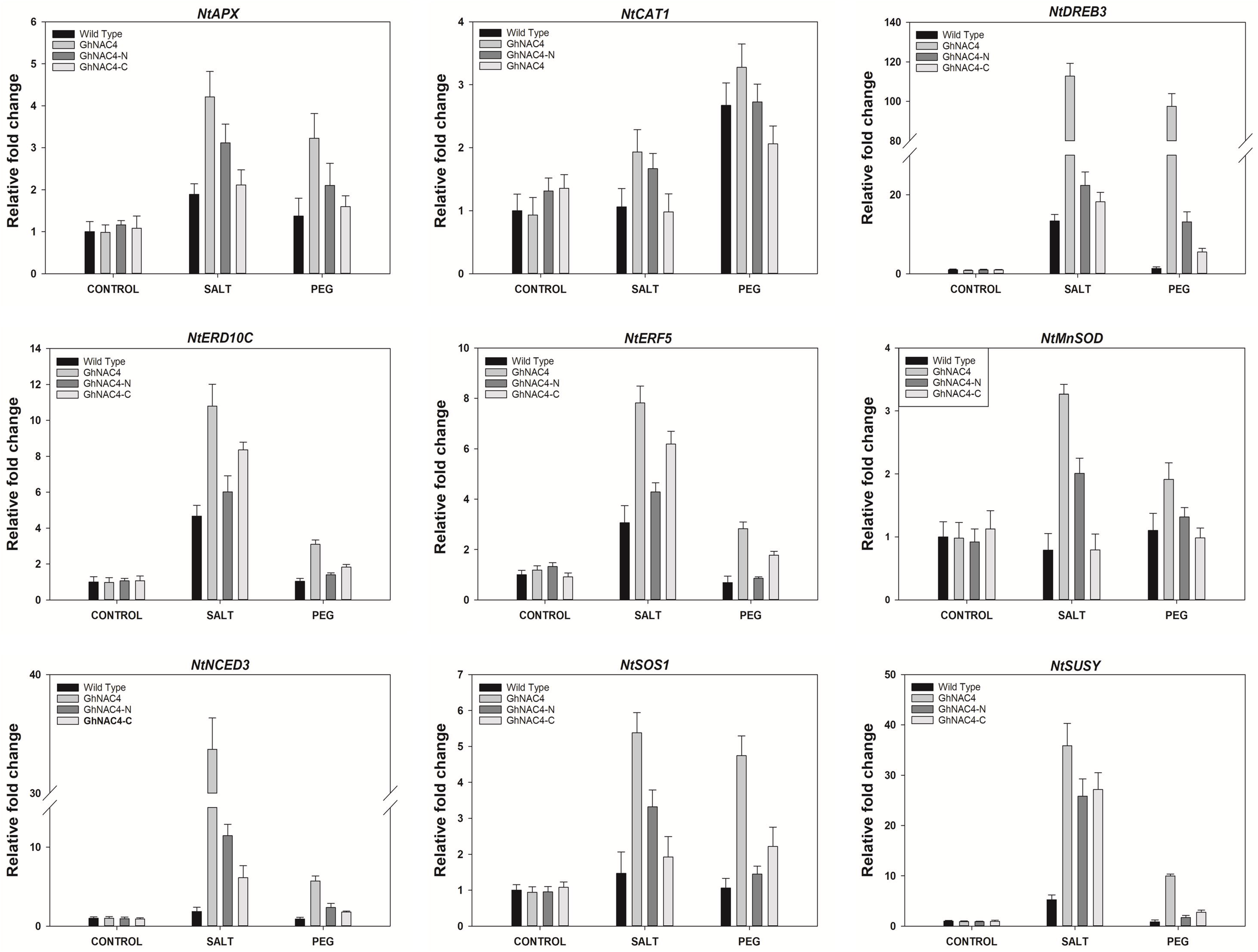
Expression patterns of stress responsive genes in WT, GhNAC4, GhNAC4-N and GhNAC4-C expressing tobacco transgenic plants in response to salt and drought treatment. qPCR expression analyses of *NtAPX*, *NtCAT1*, *NtERD10C*, *NtERF5*, *NtDREB3*, *NtMnSOD*, *NtNCED3*, *NtSOS1*, and *NtSUSY* transcripts after treatment with 200 mM NaCl and 15 % PEG 8000 for 24 h. Two weeks-old tobacco seedlings were used in the analyses. The transcript levels of the stress responsive genes were normalized with that of tobacco Ubiquitin, *NtUBI1*. The data are shown as the mean ± SE (n = 3)

The transcriptional regulatory role of GhNAC4 was supported by the altered expression of many ABA and stress-responsive genes. In the GhNAC4 genotype, the transcript levels of all the 9 genes were upregulated to a higher level under both treatments as compared to wild-type, GhNAC4-N and GhNAC4-C genotypes. Notably, a very high upregulation of *NtNCED3* and *NtDREB3* transcripts was observed. This suggests the importance of GhNAC4 in abiotic stress response. It is to be noted that the former gene is involved in ABA biosynthesis whereas the latter is a gene that works in ABA independent pathway.

In GhNAC4-N genotype, under both NaCl and PEG treatments, the expression levels of *NtAPX*, *NtCAT1*, *NtDREB3*, *NtMnSOD*, *NtNCED3*, and *NtSOS1* were upregulated to higher folds as compared to GhNAC4-C, while the expression levels of *NtERD10C*, *NtERF5* showed higher upregulation in GhNAC4-C genotype under both the stresses. This result suggests that both NAC and TR domains play a role in altering the expression of abiotic stress and ABA-responsive genes. Remarkably, GhNAC4 shows an additive effect of upregulation of stress-responsive genes by GhNAC4-N and GhNAC4-C genotypes and is a positive regulator of salinity and drought stress.

## Discussion

In the present study, the domains of a cotton NAC TF, GhNAC4 were investigated to better understand their structure-function relationship. Most NAC TFs have an N-terminal NAC-domain and a C-terminal TR-domain minimally. In this study, we demonstrated that the NAC-domain is largely responsible for the ABA responsiveness of GhNAC4. This conclusion was based on the results showing that GhNAC4-N transgenics exhibited hypersensitivity to exogenous ABA both during seed germination and seedling growth whereas the GhNAC4-C genotype transgenics were insensitive.

### Stomatal movement and water loss

It is well known that the stomatal movement is largely controlled by ABA and the measurement of the stomatal aperture is a useful indicator of altered ABA sensitivity. Treatment of epidermal peels with ABA resulted in enhanced closure of stomatal aperture in GhNAC4-N genotype as compared to GhNAC4-C genotype that exhibited insensitivity towards exogenously applied ABA (as observed by reduced stomatal closure). Manipulation of genes that modulate stomatal movements has resulted in alteration of stomatal aperture. Members of TF family such as R2R3MYB, ERF/AP2, NF-YA and NAC are known to alter stomatal movements (Cominelli et al. 2010). For instance, *SNAC1*, a stress-responsive NAC TF from rice is predominantly induced in guard cells under drought conditions. Transgenic rice over-expressing SNAC1 showed enhanced ABA sensitivity and had more rapid stomatal closure under both control and drought conditions as compared to WT plants (Hu et al. 2006). Interestingly, both GhNAC4-N and GhNAC4-C genotypes demonstrated reduced aperture size under normal conditions as compared to wild-type and GhNAC4 genotype. Mutants with altered sensitivity or deficient in ABA also exhibit altered leaf water loss condition. In our experiments, we observed that GhNAC4-C genotype had faster rate of water loss as compared to the plants of GhNAC4-N genotype corroborating the insensitivity of GhNAC4-C genotype plants to externally applied ABA. Furthermore, the reduced transpiration rate of full length GhNAC4-overexpressing plants and its strong promoter activity in guard cells (Fig. 3 Trishla et al. 2020) suggest the active role of GhNAC4 in stomatal closure.

### GhNAC4 expression and stress adaptation

In this study, we also demonstrated that GhNAC4 is involved in the adaptation to salinity and drought stress. This conclusion was based on the observation that GhNAC4-overexpressing plants showed increased accumulation of proline, reduced chlorosis and membrane damage under both salinity and drought stress treatments. GhNAC4 plants also had better growth performance as measured by germination rate and seedling growth under stress conditions. Considering the role of GhNAC4 as a TF, it is likely to play a role in regulating gene expression during abiotic stress. Indeed, quite a few stress and ABA-responsive genes (*NtNCED3*, *NtDREB3*, *NtSOS1*, *NtERD10C*, *NtERF5*, *NtAPX*, *NtCAT1*, *NtMnSOD*, and *NtSUSY*) were highly upregulated by GhNAC4 under salt and drought stress conditions. Upregulation of stress-responsive genes suggests that GhNAC4 may act as a transcriptional regulator for the downstream modulators of stress response, thereby further suggesting the importance of GhNAC4 in abiotic stress response. It is possible that GhNAC4 transcriptionally activates the stress-responsive genes by binding to the NAC recognition sequence (NACRS) in their promoter regions. We observed the presence of several NACRS in the 1 kb promoter region of all the nine genes (Supplementary Table 3). This suggests that GhNAC4 may act as a possible direct upstream regulator of abiotic stress responsive genes.

Interestingly, a very high upregulation of *NtNCED3* and *NtDREB3* transcripts was observed in GhNAC4-overexpressing plants under salinity and drought stress. Higher levels of NCED3 (9-*cis*-epoxycarotenoid dioxygenase 3), an ABA biosynthetic enzyme (Iuchi et al. 2001) points towards increased levels of endogenous ABA in GhNAC4-overexpressing plants under abiotic stress treatment. This suggests that GhNAC4 is able to directly or indirectly modulate the biosynthesis of ABA. In contrast, DREB3 (dehydration responsive element-binding protein 3) is an AP2 domain containing TF known to regulate expression of several stress-inducible gene in an ABA-independent manner (Lata and Prasad 2011). High induction of both *NtNCED3* and *NtDREB3* transcripts in GhNAC4-overexpressing plants under stress treatment suggests that GhNAC4 is likely to be involved in both ABA-dependent and independent pathways for stress signalling. This is partially supported by the fact that the promoter region of GhNAC4 contains several copies of ABA-response element (ABRE, binding site for ABRE binding proteins) and dehydration-response element (DRE, binding site for DREBs) *cis*-acting elements (Supplementary Table 2, Trishla et al. 2020). In response to abiotic stress, ABRE and DRE are involved in regulation of gene expression in ABA-dependent and independent manner, respectively (Shinozaki and Yamaguchi-Shinozaki 2007).

We demonstrated that the stress tolerance ability of GhNAC4 is associated with both the domains, NAC- and TR. Tobacco transgenics expressing both GhNAC4-N and GhNAC4-C constructs independently were tolerant to salt and drought treatments as compared to wild-type. However, GhNAC4-N genotype plants were more tolerant compared to GhNAC4-C expressing plants. GhNAC4-N genotype plants showed reduced chlorosis and lipid peroxidation and a higher proline content under abiotic stress. Also GhNAC4-N plants had better growth performance under salinity and drought stress conditions.

### GhNAC4 domains and associated upregulation of stress related genes

The transcripts of ABA and stress-responsive genes were also upregulated in GhNAC4-N and GhNAC4-C expressing tobacco transgenic plants. A higher level of expression of *NtNCED3* was observed in GhNAC4-N plants as compared to GhNAC4-C plants. This suggests the enhanced *de novo* ABA biosynthesis in GhNAC4-N plants. This also supports the notion that NAC-domain of GhNAC4 is largely responsible for the ABA-mediated signaling. Also it can possibly explain the hypersensitivity of GhNAC4-N plants to exogenous application of ABA. The SOS1 (Salt Overly Sensitive 1) encodes a plasma membrane Na^+^/H^+^ antiporter and plays a critical role in controlling the transport of Na^+^ from root to shoot, thus being essential for maintaining Na^+^ and K^+^ homeostasis. Increased expression of *SOS1* is reported to improve salt tolerance in *Arabidopsis* (Shi et al. 2003). A higher level of induction of *NtSOS1* transcripts was observed in GhNAC4-N plants as compared to GhNAC4-C plants. Salinity and drought treatments impose osmotic stress, which leads to the production of reactive oxygen species (ROS). It is critical to scavenge and decrease ROS levels to maintain cellular homeostasis (Krasensky and Jonak 2012). There is a positive correlation between abiotic stress tolerance and activity of ROS scavenging antioxidant enzymes (Wang et al. 2009). Again, a level of higher induction of ROS-scavenging genes such as catalase 1 (*NtCAT1*), ascorbate peroxidase (*NtAPX*), and superoxide dismutase (*NtMnSOD*) was observed in GhNAC4-N as compared to GhNAC4-C genotype. This can presumably explain the reduced extent of lipid peroxidation and increased stress tolerance capabilities of GhNAC4-N genotype.

Transcripts of early response to dehydration 10C (*NtERD10C*) and ethylene response factor 5 (*NtERF5*) were upregulated to a higher extent in GhNAC4-C genotype as compared to GhNAC4-N genotype under both salinity and drought stress treatments. ERD10C is an osmo-protectant hydrophilic protein that is known to play a critical role in maintaining cell homeostasis (Siqueira and Gomes 2013). Higher induction of ERD10C suggests that the plants can be better equipped with protecting macromolecules and stabilizing labile enzymes (Chakrabortee et al. 2007). ERF5 plays a regulatory role in hormone cross-talk and redox signaling under environmental stress (Pan et al. 2012). This suggests that the C-terminal TR-domain is a transcriptional activator and does contribute to the enhanced stress tolerance capabilities of GhNAC4. The functional specificity of both the NAC and TR-domains may be explained by their ability to interact with other proteins of the transcriptional complex.

### Functional significance of individual domains of GhNAC4

A characteristic feature of NAC TFs is that they can form homo- and heterodimers. The NAC-domain contains highly conserved Arginine and Glutamate residues at positions 19 and 26 respectively, which form the two prominent salt bridges essential for dimer formation (Olsen et al. 2005b). Consistent with this observation, the conserved amino acids were present in the GhNAC4 sequence and GhNAC4 also forms dimers in yeast. We observed that the N-terminal NAC-domain is essential for dimer formation and alteration of the conserved amino acids abolished the dimerization.

Another feature of NAC TFs is that they act as transcriptional activators or repressors. The C-terminal TR-domain have a high degree of intrinsic disorder essential for transcriptional regulation (Jensen et al. 2010). Consistent with this observation, the C-terminal TR-domain is essential for the transcriptional activation of GhNAC4 and interestingly had a higher degree of β-galactosidase activity as compared to the full-length protein. The intrinsically disordered regions generally contain a higher proportion of charged and polar amino acids. These regions are associated with low mean hydrophobicity and high net charge (Uversky et al. 2000). A similar pattern was observed in GhNAC4, where the two C-terminal regions (D1 and D2) necessary for the transcription activation property are rich in polar amino acids such as serine, glutamine and threonine.

In conclusion, this study elucidates that GhNAC4 can form homo-dimers and is a transcriptional activator. The significantly enhanced salinity and drought tolerance of GhNAC4 is associated with both the N-terminal NAC-domain and the C-terminal TR-domain. The ABA-responsiveness of GhNAC4 is largely associated with the NAC-domain.

## Acknowledgements

The authors would like to acknowledge Bharath P. and Dr. Krishnaveni Mishra for their help with stomatal bioassay and yeast assay respectively. VST would like to acknowledge Vikas Kumar Jain and Dr. Aswini Vetcha for the critical reading of the manuscript. The authors are grateful to the Head, Department of Plant Sciences, University of Hyderabad for the facilities under various umbrella programs like DST-FIST and UGC SAP-DRS.

## Author contributions

VST and PBK conceived and designed the experiments. VST performed the experiments and analyzed the data. VST and PBK wrote the manuscript.

## Data availability

All the data presented in the present manuscript are freely available to the interested researchers and would be made available upon request to the corresponding author – VST.

## Compliance with ethical standards

## Ethical approval

This investigation does not necessitate the use of animals and hence, does not need the approval from the Animal Ethics Committee.

## Consent to participate

The present manuscript is the result of the participation of all the authors listed in it.

## Consent for publication

The authors of the manuscript hereby give their consent for publication of the manuscript in PMB.

