## Supplementaary Figure 1 for "Structure-function relationship of *Gossypium hirsutum* NAC transcription factor, GhNAC4 with regard to ABA and abiotic stress responses"

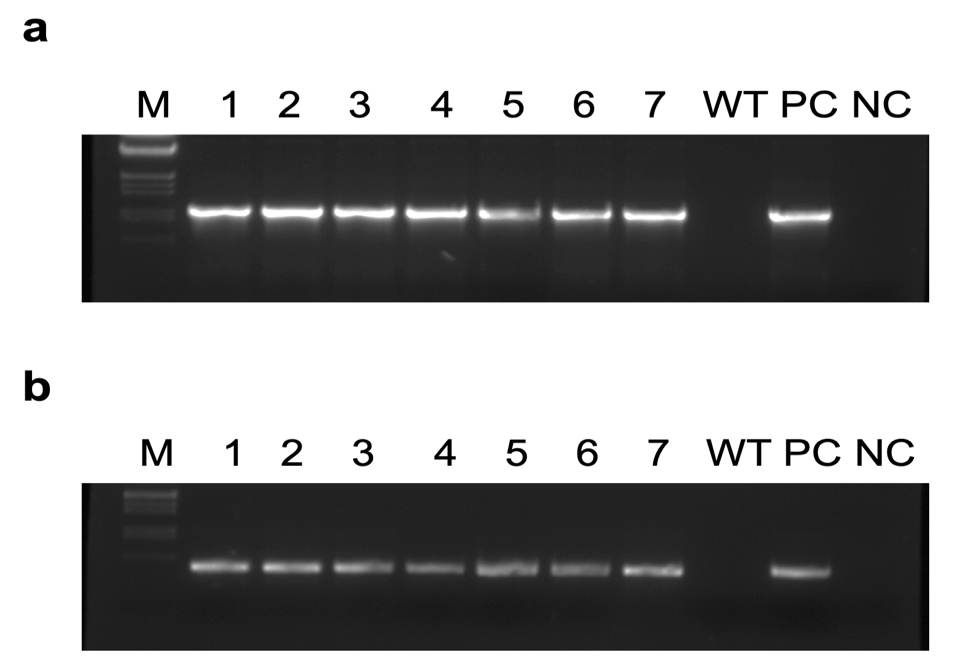

**Supplementary Figure S1**

Confirmation of T_0_ putative transgenic plants by PCR using tobacco genomic DNA. A representative gel picture showing **a** 1041 bp amplified product of GhNAC4 cds and **b** 739 bp amplified product of *nptII*. ‘M’ represents λ *EcoR*I*/Hin*dIII DNA ladder. ‘PC’ represents positive control for the PCR using plasmid (pCAMBIA2300::GhNAC4) as a template. ‘WT’ represents negative control for the PCR using DNA from untransformed plant. ‘NC’ represents no template control

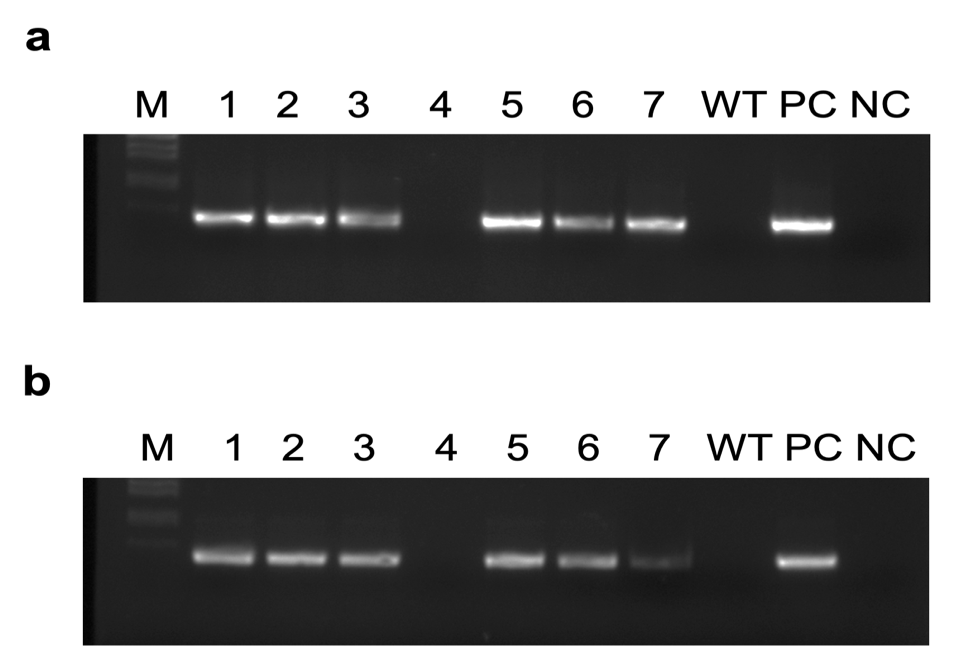

**Supplementary Figure S2**

Confirmation of T_0_ putative transgenic plants by PCR using tobacco genomic DNA. A representative gel picture showing **a** 417 bp amplified product of N-terminal NAC-domain of GhNAC4 and **b** 739 bp amplified product of *nptII*. ‘M’ represents λ *Eco*RI*/Hin*dIII DNA ladder. ‘PC’ represents positive control for the PCR using plasmid (pCAMBIA2300::GhNAC4-N) as a template. ‘WT’ represents negative control for the PCR using DNA from untransformed plant. ‘NC’ represents no template control

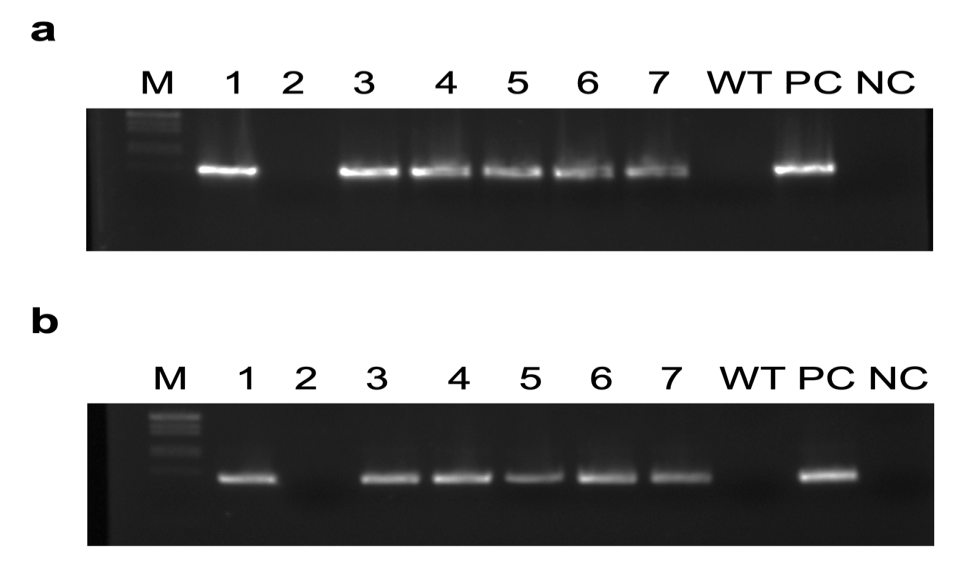

**Supplementary Figure S3**

Confirmation of T_0_ putative transgenic plants by PCR using tobacco genomic DNA. A representative gel picture showing **a** 624 bp amplified product of C-terminal transcriptional regulatory domain of GhNAC4 and **b** 739 bp amplified product of *nptII*. ‘M’ represents λ *Eco*RI/*Hin*dIII DNA ladder. ‘PC’ represents positive control for the PCR using plasmid (pCAMBIA2300::GhNAC4-C) as a template. ‘WT’ represents negative control for the PCR using DNA from untransformed plant. ‘NC’ represents no template control
