## Supplementary Table 1 for "Structure-function relationship of *Gossypium hirsutum* NAC transcription factor, GhNAC4 with regard to ABA and abiotic stress responses"

**Supplementary Table 1.** Primers used for cloning of GhNAC4 full length, NAC-domain (GhNAC4-N) and Transcriptional regulatory domain (GhNAC4-C)

| **Gene** | **Primer name** | **Primer sequence** |
| --- | --- | --- |
| *GhNAC4*  full length  (80-1120) | GhNAC4- pRT100-F | 5’ ATCTCGAGATGGGAGTGCCGGAAACTG 3’  *Xho*I |
|  | GhNAC4- pRT100-R | 5’ GCCGGATCCTTATTGTCTAAACCCAAATCC 3’  *Bam*HI |
|  | GhNAC4-pEGAD-F | 5’ ATTCCCGGGATGGGAGTGCCGGAAAC 3’  *Sma*I |
|  | GhNAC4-pEGAD-R | 5’ GCCGGATCCTTATTGTCTAAACCCAAATCC 3’  *Bam*HI |
|  | GhNAC4-yeast-F | 5’ ATTCCCGGGATGGGAGTGCCGGAAAC 3’  *Sma*I |
|  | GhNAC4-yeast-R | 5’ GCCGGATCCTTATTGTCTAAACCCAAATCC 3’  *Bam*HI |
| GhNAC4-N  (80-496) | GhNAC4-N-pRT100-F | 5’ ATTCCCGGGATGGGAGTGCCGGAAAC 3’  *Sma*I |
|  | GhNAC4-N- pRT100-R | 5’ATAGGATCCTTATCGATATTCATGCATAATC 3’  *Bam*HI |
|  | GhNAC4-N-yeast-F | 5’ ATTCCCGGGATGGGAGTGCCGGAAAC 3’  *Sma*I |
|  | GhNAC4-N-yeast-R | 5’ATAGGATCCTTATCGATATTCATGCATAATC 3’  *Bam*HI |
| GhNAC4-C  (80-496) | GhNAC4-C-pRT100-F | 5’ ATTCCCGGGATTGAAACTTCTCGTAAAAG 3’  *Sma*I |
|  | GhNAC4-C- pRT100-R | 5’ GCCGGATCCTTATTGTCTAAACCCAAATCC 3’  *Bam*HI |
|  | GhNAC4-C-yeast-F | 5’ ATTCCCGGGATTGAAACTTCTCGTAAAAG 3’  *Sma*I |
|  | GhNAC4-C-yeast-R | 5’ GCCGGATCCTTATTGTCTAAACCCAAATCC 3’  *Bam*HI  BamHI |
| *nptII* | nptII-F | AGATGGATTGCACGCAGGTTCTC |
|  | nptII-R | ATCGGGAGCGGCGATACCGTA |
