## Supplementary Table 2 for "Structure-function relationship of *Gossypium hirsutum* NAC transcription factor, GhNAC4 with regard to ABA and abiotic stress responses"

**Supplementary table 2.** Primers used for expression analysis of stress-responsive genes

| **Gene** | **Primer name** | **Primer sequence** |
| --- | --- | --- |
| *NtAPX*  (U15933.1) | NtAPX-F | 5’ GTTTGGGCTTTTCTCCTCGAC 3’ |
|  | NtAPX-R | 5’ GGAGCATAAGAGGAGCGCAA 3’ |
| *NtCAT1*  (U93244.1) | NtCAT1-F | 5’ GGCCGCTACAACTCTCTCTTT 3’ |
|  | NtCAT1-R | 5’ ACAGGACCTCTTGCACCAAC 3’ |
| *NtERD10C*  (AB049337.1) | NtERD10C-F | 5’ AAAGCCAACTCATGCCCAAG 3’ |
|  | NtERD10C-R | 5’ AGAGCTGCTACTTGATCGATGG 3’ |
| *NtERF5*  (AY655738.1) | NtERF5-F | 5’ GGATTGTCTCCTGCTGCTGT 3’ |
|  | NtERF5-R | 5’ GCTCTTCTAATAACTCAGCACCC 3’ |
| *NtDREB3*  (EU727157.1) | NtDREB3-F | 5’ ATGGCTTGGCACTTTCCCTT 3’ |
|  | NtDREB3-R | 5’ ATATTCTTGGCGTCGGAGGA 3’ |
| *NtMnSOD*  (AB093097.1) | NtMnSOD-F | 5’ TCCCCTACGACTATGGAGCA 3’ |
|  | NtMnSOD-R | 5’ CGGTATGCAATTTGGCGACG 3’ |
| *NtNCED3*  (JX101472.1) | NtNCED3-F | 5’ TGTCTGAAATGATCCGGGGC 3’ |
|  | NtNCED3-R | 5’ AGTTTCCGGCTCTTCCCAAG 3’ |
| *NtSOS1*  (XM_009789739.1) | NtSOS1-F | 5’ CAAATGTTATCCCCCGAAAGC 3’ |
|  | NtSOS1-R | 5’ CGGAGAACCTGAGGAAATGTGA 3’ |
| *NtSUSY*  (AB055497.1) | NtSUSY-F | 5’ CACGGATATTTCGCCCAGGA 3’ |
|  | NtSUSY-R | 5’ GCAGCAGCCGAGTAGCAATA 3’ |
| *NtUBI1*  (U66264.1) | NtUBI1-F | 5’ GAGTCAACCCGTCACCTTGT 3’ |
|  | NtUBI1-R | 5’ ACATCTTTGAGACCTCAGTAG ACA 3’ |
