## Supplementary Table 3 for "Structure-function relationship of *Gossypium hirsutum* NAC transcription factor, GhNAC4 with regard to ABA and abiotic stress responses"

**Supplementary Table 3.** Putative NAC TF recognition sequence (NACRS) in the promoters of tobacco stress-responsive genes identified using PlantPAN 3.0 ([http://PlantPAN.itps.ncku.edu.tw](http://plantpan.itps.ncku.edu.tw/)). One kb upstream region was used for analysis. Similarity score > 0.9 used for prediction

| Gene | GenBank Accession no. | NACRS |
| --- | --- | --- |
| *NtDREB3* | EU727157.1 | 4 |
| *NtCAT1* | U93244.1 | 12 |
| *NtAPX* | U15933.1 | 12 |
| *NtERD10C* | AB049337.1 | 7 |
| *NtERF5* | AY655738.1 | 11 |
| *NtMnSOD* | AB093097.1 | 2 |
| *NtNCED3* | JX101472.1 | 10 |
| *NtSOS1* | XM_009789739.1 | 9 |
| *NtSUSY* | AB055497.1 | 12 |
